# A primary microcephaly-associated *sas-6* mutation perturbs centrosome duplication, dendrite morphogenesis, and ciliogenesis in *Caenorhabditis elegans*

**DOI:** 10.1101/2022.11.25.518003

**Authors:** Mary Bergwell, Amy Smith, Ellie Smith, Carter Dierlam, Ramon Duran, Erin Haastrup, Rebekah Napier-Jameson, Rory Seidel, William Potter, Adam Norris, Jyoti Iyer

## Abstract

The human SASS6(I62T) missense mutation has been linked with the incidence of primary microcephaly in a Pakistani family, although the mechanisms by which this mutation causes disease remain unclear. The SASS6(I62T) mutation corresponds to SAS-6(L69T) in *C. elegans*. Given that SAS-6 is highly conserved, we modeled this mutation in *C. elegans* and examined *sas-6(L69T)* effect on centrosome duplication, ciliogenesis and dendrite morphogenesis. Our studies revealed that all the above processes are perturbed by the *sas-6(L69T)* mutation. Specifically, *C. elegans* carrying the *sas-6(L69T)* mutation exhibit an increased failure of centrosome duplication in a sensitized genetic background. Further, worms carrying this mutation also display shortened phasmid cilia, an abnormal phasmid cilia morphology, shorter phasmid dendrites and chemotaxis defects. Our data show that the centrosome duplication defects caused by this mutation are only uncovered in a sensitized genetic background, indicating that these defects are mild. However, the ciliogenesis and dendritic defects caused by this mutation are evident in an otherwise wild-type background, indicating that they are stronger defects. Thus, our studies shed light on the novel mechanisms by which the *sas-6(L69T)* mutation could contribute to the incidence of primary microcephaly in humans.

## Introduction

Autosomal recessive primary microcephaly (MCPH) is a rare disease that causes a reduced brain size in infants along with a host of other physical and intellectual disabilities (Zaqout et al. 2017). According to the OMIM database, thirty microcephaly loci named MCPH1-MCPH30 have been identified thus far (https://www.omim.org/entry/251200). The current consensus in the field is that most mutations linked to the incidence of MCPH deplete the pool of neural progenitor cells during brain development. This results in fewer neurons being produced during embryonic development, thereby leading to a smaller brain size (Phan and Holland 2021). Although some MCPH-associated mutations have been previously modeled in *in vivo* model systems to understand how MCPH-associated mutations contribute to the disease pathology, there are still many MCPH-linked mutations that remain uninvestigated (Phan and Holland 2021). One such MCPH-associated mutation is a mutation within the *SASS6* gene (*sas-6* in *C. elegans*) (Khan et al. 2014). In 2014, a study by Khan et al. identified a leucine to threonine missense mutation in the human *SASS6* gene to be associated with the incidence of MCPH in a consanguineous Pakistani family (Khan et al. 2014).

SAS-6 protein function is conserved amongst several species including the multicellular eukaryotic nematode *C. elegans* and humans (Dammermann et al. 2004, Leidel et al. 2005). The *sas-6* gene was first identified to regulate centrosome duplication in *C. elegans* (Dammermann et al. 2004, Leidel et al. 2005). Centrosomes consist of a pair of cylindrical centrioles: a mother and a daughter centriole that are embedded in pericentriolar material. Major functions of these organelles involve organizing a mitotic spindle and aiding in cilia assembly (Nigg and Holland, 2018). Proper cell division requires a precise duplication of centrosomes. There are two levels of regulation of centrosome duplication. First, it is critical that centrioles duplicate only once per cell cycle, and second, it is imperative that each mother centriole gives rise to a single daughter centriole. Defects in centrosome duplication can impair proper chromosome segregation, leading to genome instability and aneuploidy. Indeed, increasing centrosome numbers by elevating the expression levels of the master centrosome duplication kinase Plk4 (ZYG-1 in *C. elegans*) is sufficient to induce tumorigenesis in mice (Levine et al. 2017). ZYG-1 and SAS-6 both play a critical role in mediating centrosome duplication (O’Connell et al. 2001, Dammermann et al. 2004, Leidel et al. 2005). SAS-6 is recruited to the mother centriole through its direct interaction with ZYG-1 and with another centriolar protein SAS-5 (Lettman et al. 2013). These interactions are essential to initiate centriole assembly.

In addition to regulating centrosome biogenesis, SAS-6 function is also essential for cilia assembly in *C. elegans* (Li et al. 2017). Cilia are microtubule-based organelles that are critical for cell signaling and development (Drummond 2012). Sensory cilia are the sole type of cilia that are present in *C. elegans.* These cilia are non-motile in nature and they play an essential role in sensing the extracellular environment (Inglis et al. 2007). There are a total of sixty ciliated sensory neurons in *C. elegans* hermaphrodites. The cilia are present at the dendritic ends of the sensory neurons (Inglis et al. 2007). Although a previous study has shown that SAS-6 function is critical for proper ciliogenesis (Li et al. 2017), the mechanism by which SAS-6 promotes ciliogenesis is unknown. It has been especially difficult to study the role of SAS-6 in ciliogenesis because *sas-6/SASS6* is an essential gene (Kamath et al. 2003, Blomen et al. 2015). Due to the important role of SAS-6 in regulating centrosome duplication, most mutations in *sas-6* yield embryonic lethality or sterility (Dammermann et al. 2004, Leidel et al. 2005, O’Rourke et al. 2011, Song et al. 2011, The C. elegans Deletion Mutant Consortium et al. 2012, Lettman et al. 2013).

Ciliogenesis defects have been proposed as an important underlying cause of microcephaly (Alcantara et al. 2014, Gabriel et al. 2016, Broix et al. 2018, Wambach et al. 2018, Zhang et al. 2019, Ding et al. 2019, Sohayeb et al. 2020, Farooq et al. 2020). However, the effect of this *SASS6* mutation on ciliogenesis has never been investigated. The effect of the MCPH-associated *SASS6* mutation on centrosome duplication in its endogenous context has also not been determined thus far. It is important to note that SASS6 levels in the cell are tightly regulated. An increase or a decrease in SASS6/SAS-6 levels causes either centrosome overduplication or a failure of centrosome duplication, respectively (Dammermann et al. 2004, Leidel et al. 2005). Therefore, it is critical to model the MCPH-associated *SASS6* mutation in an endogenous context in a multicellular eukaryotic animal model. However, to date such a study has not been reported.

In this study, we used CRISPR/Cas9 genome editing to re-create the MCPH-associated *sas-6(L69T)* mutation in *C. elegans*. Our *in vivo* studies in the worm indicate that the *sas-6(L69T)* mutation does not result in embryonic lethality when this is the sole mutation in *sas-6* in an otherwise wild-type background. However, introducing this mutation into a tight temperature-sensitive *zyg-1(it25)* strain that is partially compromised for centrosome duplication results in a robust increase in embryonic lethality at the semi-permissive temperature. These data indicate that this MCPH-associated mutation mildly perturbs centrosome duplication in *C. elegans.* Surprisingly, while the centrosome duplication defects were only observed in a sensitized genetic background, ciliogenesis and dendritic defects were observed even in a wild-type background in *C. elegans* carrying the *sas-6(L69T)* mutation. Specifically, our studies reveal that the *sas-6(L69T)* mutation causes a reduction of the phasmid cilia and phasmid dendrite lengths and results in an abnormal phasmid cilia morphology. Thus, our studies indicate that the MCPH-associated *sas-6(L69T)* mutation affects ciliogenesis and dendrite morphogenesis more severely than centrosome duplication in *C. elegans.* In conclusion, our studies have uncovered a novel role of the MCPH-associated *sas-6(L69T)* mutation in regulating ciliogenesis and dendrite morphogenesis in the worm.

## Materials and Methods

### *C. elegans* maintenance

All *C. elegans* strains were grown at 20°C on OP50 *E. coli* bacteria seeded onto MYOB agar plates, unless otherwise indicated (as in the case of different assays). **Table S1** lists all the *C. elegans* strains that have been generated and used in this study.

### CRISPR/Cas9 genome editing

Assembled ribonucleoprotein complexes were microinjected into *C. elegans* gonads to perform CRISPR/Cas9 genome editing as described previously (Paix et al. 2015, Paix et al. 2016, Iyer et al. 2019, Smith et al. 2020). Homozygosity of the *sas-6(L69T)* edit was confirmed by PCR followed by restriction digestion with EcoRI and by DNA sequencing. Note that the EcoRI restriction site was introduced by silent mutagenesis of the *sas-6* gene sequence. Therefore, it is not expected to alter the final sequence of the SAS-6 protein. **Table S2** lists the sequences of the screening primers, guide RNAs and repair templates that were used to make the CRISPR strains that have been used in this study.

### Brood count and embryonic viability assays

All brood count assays and embryonic viability assays were performed as described previously (Xie et al. 2022). For both brood count as well as embryonic viability assays, L4 stage *C. elegans* were plated at the respective indicated temperatures. For brood count assays, all the progeny that were produced by each single worm over the worm’s reproductive lifespan were counted and quantified. For embryonic viability assays, the percentage of dead and live progeny produced by each worm was determined by counting. The percentage of viable progeny was then computed by dividing the number of live progeny by the total number of progeny and by multiplying this ratio by 100.

### Confocal live imaging of centrosome duplication

To monitor centrosome duplication in *C. elegans* embryos, live imaging was performed using an Olympus Spinning-Disk confocal microscope mounted with a Yokagawa scanhead. The cellSens software (Olympus America Inc., Waltham, MA) was used for image acquisition and processing. To prepare the *C. elegans* strains for imaging, L4 stage *C. elegans* were plated at 21.8°C overnight. The next day, gravid adult worms were picked into 8 to 12 µl of egg buffer (118 mM NaCl, 47 mM KCl, 2 mM CaCl_2_, 2 mM MgCl_2_, 5 mM HEPES pH 7.4) on a square glass coverslip and dissected near the vulva using a needle to release embryos. The coverslip was carefully mounted on a 2% agarose pad (2% agarose LE in M9 buffer). The edges of the coverslip were sealed with molten Vaseline using a paintbrush. These slides were subjected to time-lapse imaging by taking 1.25 µm Z-stacks over an interval of approximately 132 seconds.

Maximum intensity Z-projections were computed using the cellSens software (Olympus America Inc.) and converted into an AVI format for making videos. The brightness and contrast were adjusted for the control and *sas-6(L69T)* videos for better non-quantitative video visualization.

### Genetic crosses

To introduce the *sas-6(L69T)* mutation into the *zyg-1(it25)* background, first, males were generated from the *zyg-1(it25)* strain using alcohol treatment. Specifically, L4 stage hermaphrodites from the *zyg-1(it25)* strain were incubated with 10% ethanol in M9 buffer for 30 minutes, washed thrice with M9 buffer and allowed to recover for an hour. After recovery, the worms were plated onto MYOB plates and incubated at 20°C until males appeared. Following this, *zyg-1(it25)* homozygous males were mated with *sas-6(L69T)* L4 hermaphrodites. The F1 progeny from this cross were screened for heterozygosity of the *sas-6(L69T)* mutation using PCR and restriction digestion. Cross progeny were identified and screened for homozygosity of both edits using PCR and restriction digestion (for homozygosity screening of the *sas-6(L69T)* mutation) and 24°C temperature shifts (for homozygosity screening of the *zyg-1(it25)* mutation). Homozygosity of fluorescent markers was confirmed by confocal fluorescence imaging.

For the cilia imaging experiments, PY6100 males *(oyIs59[osm-6p::gfp] III)* were crossed with hermaphrodites from the IYR001 (*sas-6(luv1[sas-6::ha]) IV*) and IYR002 (*sas-6(luv2[sas-6[L69T])) IV*) strains to produce strains expressing either SAS-6::HA (control) or SAS-6(L69T)::HA along with a pan-ciliary GFP marker. These strains were named IYR026 and IYR027, respectively. F1 progeny were screened for heterozygosity at both loci by screening F1 hermaphrodites for the presence of GFP fluorescence in the phasmid neurons. F2 homozygotes for *sas-6::ha* were identified as described previously (Bergwell et al. 2019) by PCR and digestion by NdeI followed by agarose gel electrophoresis. F2 homozygotes for *sas-6(L69T)::ha* were identified by PCR and digestion by EcoRI followed by agarose gel electrophoresis.

### Modeling SAS-6(L69T) using AlphaFold2

The N-terminal coiled-coil dimer construct of *C. elegans* centriolar protein SAS-6 was crystallized by Hilbert et al. 2013. The ceSAS-constructs required for crystallization involved the removal of the α2-β5 loop (Δ103-130) which was reported to not affect the N-terminal domain structure within a C_α_ rmsds of ∼0.3 Angstroms. The crystallographic data was deposited in the Research Collaboration for Structural Bioinformatics databank under accession number PDB ID code 4G79.

The native SAS-6 structure (AF-062479-F1) available from the AlphaFold Protein Structure Database (Jumper et al. 2021; https://AlphaFold.ebi.ac.uk/) has leucine at position 69 at the end of the β_4_ sheet just prior to the asparagine-asparagine which initiates the α_1_ helix occurring at positions 70 to 82 (using the Hilbert et al. 2013 author numbering system). It should be noted that the residue numbering system reported for ceSAS-6 in Uniprot (062479) has two additional amino acids (glycine-serine) at the N-terminus and thus the sequence is offset and the α_1_ helix starts at position 72 in this numbering system. The reported per-residue confidence score (pLDDT) for residue leucine 69 was 92.37 suggesting that the AlphaFold model confidence level was very high for this residue.

The SAS-6(L69T) mutant protein structure containing the single leucine to threonine substitution was modeled by loading the modified sequence into the Google Colaboratory (Colab) services using ColabFold: AlphaFold2 using MMseqs2 as described by Mirdita et al. 2022, and available through the AlphaFold2.ipynb v2.1.0. with the suite of public notebooks available through the deepmind/AlphaFold github. The 4G79 and the AlphaFold models were uploaded into the program ChimeraX (https://www.cgl.ucsf.edu/chimerax/) developed from the Resource for Biocomputing, Visualization, and Informatics for Visualization (RBVI), University of California, San Francisco.

The Molecular Lipophilicity Potential (MLP) is a construct that spreads atomic values out in 3D. The MLP coloring on the molecular surface is based on pyMLP (https://www.rbvi.ucsf.edu/chimerax/docs/user/commands/mlp.html) as originally developed by Broto et al. 1984 and Laguerre et al. 1997 and as modified https://www.cgl.ucsf.edu/chimera/data/mlp-sep2016/index.html and was used without any color scale modifications.

### *C. elegans* phasmid cilia and dendrite imaging

*C. elegans* phasmid cilia and dendrites were imaged as described previously (Xie et al. 2022). For the phasmid and dendrite imaging experiments, a new strain IYR040 (*oyIs59 [osm-6p::gfp] III; sas-6(luv40)*) *IV*) was created by CRISPR/Cas9 editing of the IYR026 strain. Specifically, to generate the IYR040 strain, the IYR026 strain (*oyIs59 [osm-6p::gfp] III; sas-6(luv1[sas-6::ha]) IV*) was CRISPR-edited using the same guide RNA and the repair template that was used to introduce the leucine 69 to threonine (L69T) mutation, with the only exception that the IYR040 repair template lacked the L69T mutation. As a result, the IYR040 strain is genetically identical to the IYR027 strain except that it is wild-type for SAS-6. This strain served as a control for all of our phasmid cilia and dendrite imaging experiments. For the phasmid cilia and dendrite imaging assays, L4 stage *C. elegans* from the IYR040 and IYR027 strains were grown at 16°C overnight. The next day, young adult worms were anaesthetized using 10 mM Levamisole, mounted on 2% agarose pads and imaged within 30 minutes using an Olympus IX83 Spinning Disk confocal microscope coupled with a Yokogawa X1 Scan head (Olympus America Inc.). 0.25 µm Z-slices were taken from the top to the bottom of each phasmid neuron using the cellSens software (Olympus America Inc.). All images were taken at the same laser intensity and exposure time.

### Phasmid cilia and dendrite length quantification

To quantify phasmid cilia and dendrite lengths, the raw .vsi files that were captured using the cellSens software (Olympus America Inc.) were opened in ImageJ using the OlympusImageJ Plugin. ImageJ (Abràmoff et al. 2004) was then used to convert the raw data into maximum intensity Z-projections. The scale was set to 1 pixel = 110 nm. Since the expression of GFP in the phasmid neurons varied from worm to worm, the brightness and contrast were adjusted non-uniformly to delineate the beginning and end of each phasmid cilium to ensure an accurate measurement of the phasmid cilia lengths. For the dendrite imaging, the single frame view was used in cellSens (Olympus America, Inc.) to visualize the beginning and end of each phasmid dendrite in 3D to ensure that the dendrite lengths were being properly measured. Dendrites and cilia whose beginning or end could not be delineated were excluded from our analysis. To measure cilia and dendrite lengths, a line was traced along the entire length of the cilium or the dendrite using the Segmented Lines tool in ImageJ (Abràmoff et al. 2004). The lengths of the shorter of the two phasmid dendrites and the longer of the two phasmid cilia were then measured in micrometers using ImageJ (Abràmoff et al. 2004) and have been presented in **Figures 5D** and **7C**.

### Criteria used for quantifying phasmid cilia morphology defects

Phasmid cilia that did not exhibit the canonical phasmid cilia structure with two transition zones and cilia projecting from the transition zones were quantified as having an abnormal phasmid cilia morphology. Specifically, instances where the transition zones could not be clearly distinguished from the ciliary axoneme or where we were unable to clearly identify the transition zone due to additional transition zone-like GFP aggregates being present in the vicinity of the transition zone were classified as being abnormal.

### *C. elegans* chemotaxis assay

Olfaction assays were performed as previously described in Liang et al. 2022. Briefly, *C. elegans* were maintained by standard techniques on NGM agar plates seeded with OP50 *Escherichia coli* at 20^°^C. Worms were synchronized to L4 stage. Worms were washed off the plates using M9 buffer, allowed to gravity settle, supernatant removed, and then washed a further 4 times. Unseeded NGM plates (50 mM NaCl, 1.7% agar, 0.25% peptone, 1 mM CaCl_2_, 0.013 mM Cholesterol, 1 mM MgSO_4_, 25 mM KPO_4_) were divided into four quadrants, two with solvent and two with odorant. Ethanol was used as a solvent. A 1:500 dilution of the odorant butanone was made in ethanol. The Chemotaxis Index (CI) was calculated using the formula CI = (# animals in two odorant quadrants)/(# animals in any of the four quadrants). Four plates were used per genotype per experiment. Assays were repeated 3 times. For each of the three independent trials, 569, 704 and 1552 control (*sas-6::ha*) worms respectively, and 522, 526 and 663 *sas-6(L69T)* worms respectively, were analyzed for their chemotaxis towards 1:500 butanone.

### Whole worm extracts for western blotting

Whole worm extracts for western blotting were prepared as described previously (Xie et al. 2022). Briefly, 100 gravid adults from each genotype were washed with M9 buffer twice. The M9 buffer was removed and the worm pellet was resuspended in 40 µl of 4X SDS PAGE sample buffer, boiled for 10 minutes at 95°C and stored at −30°C until further use.

### Western blotting

Western blotting was performed using the Bio-Rad wet-transfer method as described previously (Xie et al. 2022). Briefly, for western blotting of whole worm extracts, 10 to 14 µl of extracts were loaded onto each well of a 7.5% SDS-PAGE gel. The gels were run until proper band separation was achieved and the protein was transferred onto a 0.2 µm nitrocellulose membrane using the wet-transfer method (Bio-Rad Laboratories). The membranes were blocked with Odyssey Blocking Buffer (LiCOR Biosciences, Inc.), probed with anti-HA (Cell Signaling, catalog C29F4 Rabbit mAb #3724) and anti-tubulin (Santa Cruz Biotechnology, Inc., catalog sc-32293) antibodies overnight at 4°C. The membranes were washed thrice with 1X TBST (0.1% w/v Tween-20, 50 mM Tris-Cl, pH 7.6, 150 mM NaCl) and incubated with IRDye® 800CW Donkey anti-Rabbit IgG secondary antibody (LiCOR Biosciences, Inc. catalog # 926-32213) and IRDye® 680RD Goat anti-Mouse IgG secondary antibody (LiCOR Biosciences, Inc. catalog # 926-68070) for 1 hour at room temperature. The membranes were washed again thrice with 1XTBST and imaged using the LI-COR Odyssey CLx imager (LI-COR Biosciences, Inc.). The band intensities were adjusted equally for all the gel lanes for better data visualization and presentation. **Table S3** lists all the antibodies that have been used in this study.

### Western blot quantification and image presentation

The band intensities across the entire membrane were uniformly adjusted using the LiCOR Image Studio software (LI-COR Biosciences, Inc.) and saved in a .tiff format for performing quantitative analysis of the bands. Quantitative western blotting was performed using ImageJ (Abràmoff et al. 2004). Briefly, the band intensities for SAS-6::HA were determined using ImageJ and normalized with respect to the loading control Tubulin. ImageJ was also used to rotate and crop representative gel images presented in **Figure S1** and in all other fluorescence images that are presented in this paper. For all images, scale bars from the original images were cropped and grouped with their corresponding original images.

### Statistical analysis

Graphpad Prism (GraphPad Software, Inc.) was used for performing statistical analysis of the data. Unpaired two-tailed t-tests were performed to determine statistical significance of the data for the brood counts, western blot assays, phasmid cilia lengths, phasmid dendrite lengths and chemotaxis assays. The Fisher’s exact test was performed to determine if the phasmid cilia morphology defects were statistically significant. A p-value of less than 0.05 was considered to attain statistical significance.

## Results

An isoleucine to threonine substitution of SASS6 was reported to be linked with the occurrence of MCPH in a Pakistani family (Khan et al. 2014). As shown previously by Khan et al., alignment of the amino acid sequences of the SAS-6 protein of different species showed that the MCPH-associated amino acid of human SASS6 is remarkably conserved among several species including *C. elegans* (Khan et al. 2014). Notably, the *sas-6* gene was first identified in *C. elegans* and was subsequently shown to function similarly in humans (Dammermann et al. 2004, Leidel et al. 2005). Due to the high functional conservation of SAS-6, we determined that *C. elegans* would be an appropriate eukaryotic model to thoroughly investigate the consequences of this mutation. The human SASS6(I62T) mutation corresponds to SAS-6(L69T) in *C. elegans*.

### The *sas-6(L69T)* mutation was successfully introduced into *C. elegans* by CRISPR/Cas9 editing

Using CRISPR/Cas9 genome editing, we generated a leucine to threonine substitution at amino acid 69 of the *C. elegans* SAS-6 protein by editing the endogenous *sas-6* gene of *C. elegans.* This strain was named IYR002 and the corresponding allele was named *sas-6(luv2).* For ease of understanding, this strain has been referred to as the *sas-6(L69T)* strain and the corresponding MCPH-associated mutation has been referred to as the *sas-6(L69T)* mutation throughout this manuscript. The repair template used for introducing the *sas-6(L69T)* mutation carried an EcoRI restriction site **(Figure 1A)**. As shown in **Figure 1B**, upon PCR amplification and restriction digestion of the *sas-6* gene with EcoRI, several homozygous edited worms were obtained in the F2 generation (lanes with black stars). The incorporation of the *sas-6(L69T)* mutation was also confirmed by DNA sequencing.

**Figure 1.**
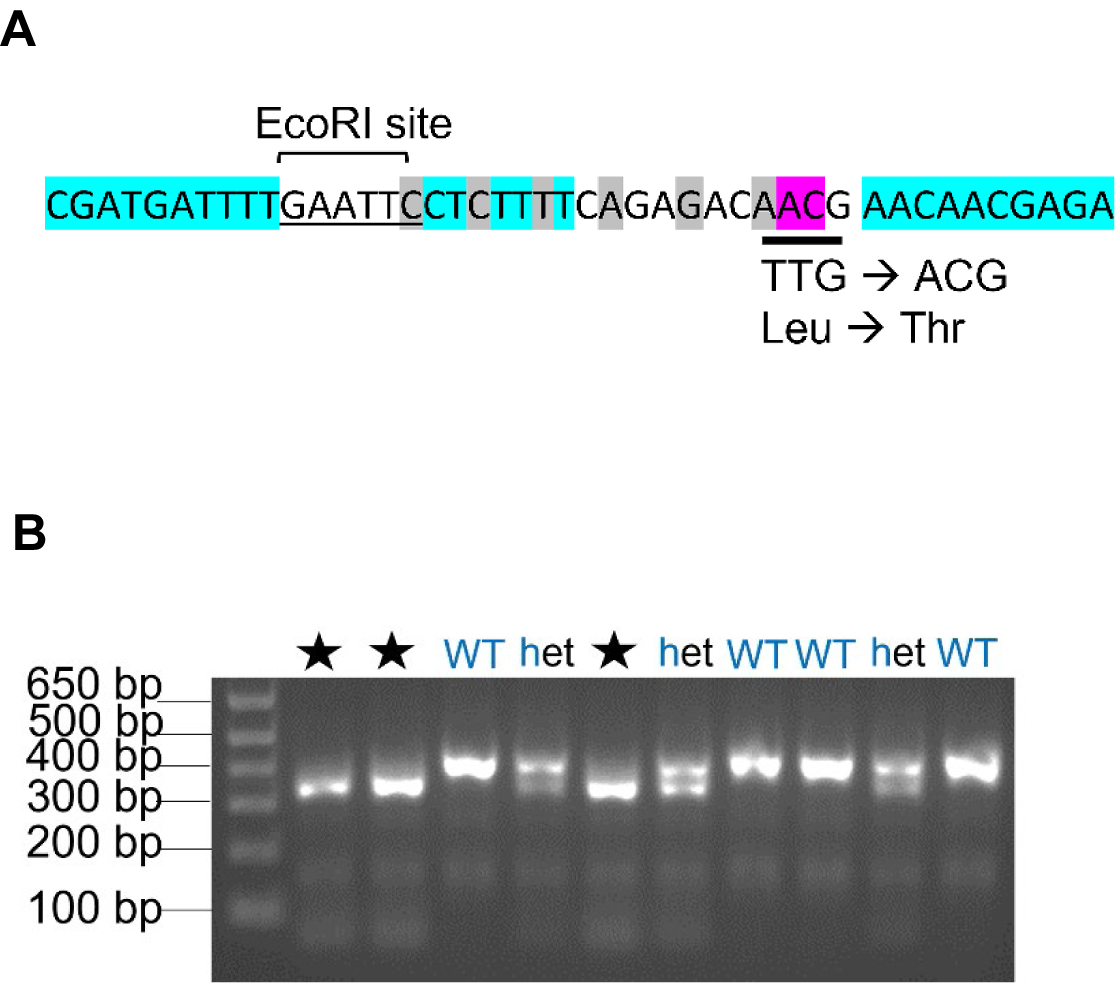
The MCPH-associated *sas-6(L69T)* mutation was successfully generated in *C. elegans* by CRISPR/Cas9 editing. A) Schematic of the CRISPR repair template used for inserting the leucine to threonine mutation at amino acid 69 of the *C. elegans* SAS-6 protein (Note: Only 10 out of 45 bases of the 5’ and 3’ homology ends of the repair template are shown). The teal highlights represent bases that are identical between the repair template and the *sas-6* gene sequence. The magenta highlight indicates the bases that were altered to introduce the leucine 69 to threonine mutation into the endogenous *sas-6* gene. The bases highlighted in gray represent silent mutations that were introduced in the repair template to prevent its cutting by the Cas9 enzyme. B) Agarose gel electrophoresis of PCR products digested with EcoRI to screen for homozygous-edited *C. elegans* carrying the *sas-6(L69T)* mutation. Black stars: Homozygous-edited worms; het: heterozygote; WT: wild-type.

### The microcephaly-associated *sas-6(L69T)* mutation does not affect brood size or embryonic viability in a wild-type background

The function of the *sas-6* gene is essential for centrosome duplication (Dammermann et al. 2004, Leidel et al. 2005). Proper centrosome duplication is required for accurate embryonic divisions in *C. elegans.* Since the *sas-6(L69T)* mutation has a pathological relevance in humans, we proposed that this mutation could impair SAS-6 function in regulating centrosome duplication, thereby leading to defective centrosome duplication in *C. elegans*. This could, in turn, affect *C. elegans* brood size and embryonic viability. Therefore, to investigate the effect of the *sas-6(L69T)* mutation on centrosome duplication, brood counts and embryonic viability assays were performed using *C. elegans* carrying this mutation. Surprisingly, the introduction of the *sas-6(L69T)* mutation did not yield any major effect on either brood size or embryonic viability at 16^°^C, 20^°^C and 25^°^C as compared with control worms (wild-type N2 worms with HA-tagged endogenous *sas-6*) **(Figures 2A and 2B)**. These data indicate that the *sas-6(L69T)* mutation does not severely perturb SAS-6 function in regulating centrosome duplication.

**Figure 2.**
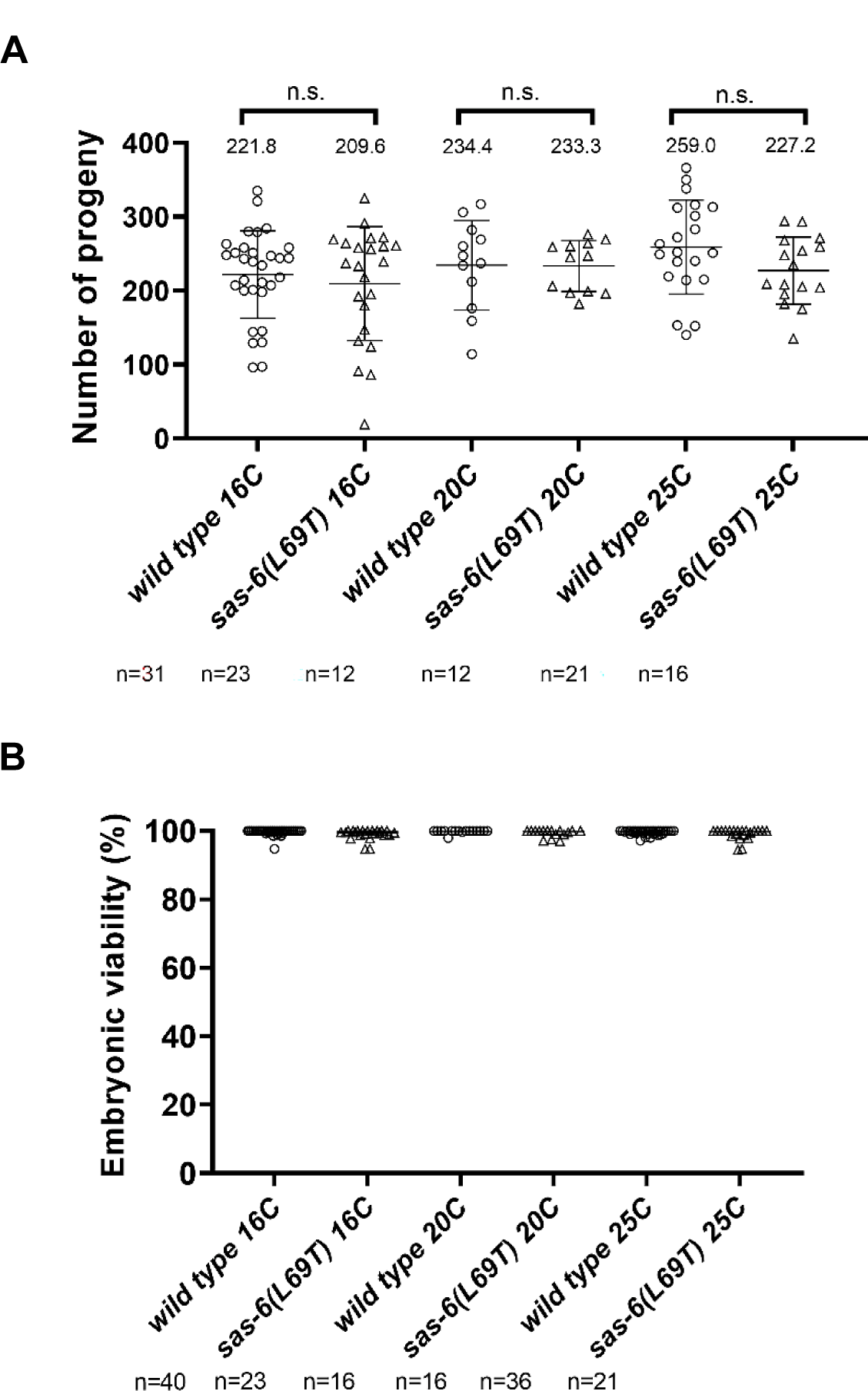
The *sas-6(L69T)* mutation does not affect *C. elegans* brood size or embryonic viability. A) Brood sizes of wild-type and *sas-6(L69T)* mutant worms were determined at temperatures of 16°C, 20°C and 25°C. Each circle or triangle represents the brood size of a single adult hermaphrodite. No statistically significant difference in brood size was observed between wild-type and *sas-6(L69T)* worms at any of the assessed temperatures. n= number of worms whose brood was analyzed for each strain. n.s: not significant. Error bars represent the standard deviation. The middle bar represents the mean. B) Embryonic viability of wild-type and *sas-6(L69T)* worms was analyzed at temperatures of 16°C, 20°C and 25°C. Each circle or triangle represents the embryonic viability of a single adult hermaphrodite. The middle line represents the median embryonic viability with a 95% confidence interval. No major change in embryonic viability was observed between wild-type and *sas-6(L69T)* worms at any of the assessed temperatures. The viability of over 2000 embryos was analyzed for each strain at every temperature.

### The *sas-6(L69T)* mutation increases centrosome duplication failures and embryonic lethality in a sensitized *zyg-1(it25)* background

ZYG-1 is a functional homolog of human Plk4 and a master kinase whose activity is critical for proper centrosome duplication (O’Connell et al. 2001, Habedanck et al. 2005, Bettencourt-Dias et al. 2005). Decreasing ZYG-1 protein levels causes a failure of centrosome duplication while elevating them causes centrosome overduplication (O’Connell et al. 2001, Peel et al. 2017, Wolf et al. 2018). The temperature-sensitive *zyg-1(it25)* allele has a proline to leucine substitution within the ZYG-1 cryptic polo box which impairs ZYG-1 function at the restrictive temperature, leading to a failure of proper centrosome duplication (Kemphues et al. 1988, Kemp et al. 2007). By controlling the temperature at which the *zyg-1(it25)* strain is grown, the rate of centrosome duplication failure in this strain can be controlled. For example, at a restrictive temperature of 24°C and above, the *zyg-1(it25)* strain produces 100% dead embryos due to a complete failure of centrosome duplication (Kemp et al. 2007). However, at lower semi-permissive temperatures (temperatures below 24°C), centrosome duplication still partially occurs in the *zyg-1(it25)* strain, yielding varying levels of embryonic viability with a temperature of 20°C yielding between 93.7% to 98.7% embryonic viability (Song et al. 2008). ZYG-1 directly interacts with SAS-6 (Lettman et al. 2013) and functions upstream of SAS-6 in the centrosome duplication pathway (Leidel et al. 2005, Delattre et al. 2006, Pelletier et al. 2006). Therefore, we questioned whether we could use the *zyg-1(it25)* allele to uncover any potential mild defects in SAS-6 function in regulating centrosome duplication that could be present in the *sas-6(L69T)* mutants. To address this, we first obtained a *zyg-1(it25)* strain expressing fluorescently-tagged centrosome (mCherry::SPD-2) and chromosome (GFP::histone) markers (OC869 from the O’Connell lab) to visualize centrosome duplication. This strain served as a control for our experiment and has been referred to as the *zyg-1(it25)* strain in **Figure 3**. Using this strain as the background strain, as shown in **Figure 3A**, we performed genetic crossing to combine the *sas-6(L69T)* and *zyg-1(it25)* mutations. This strain has been referred to as the *zyg-1(it25);sas-6(L69T)* strain in **Figure 3.** We reasoned that if the *sas-6(L69T)* mutation mildly affects SAS-6 function in regulating centrosome duplication, combining it with the *zyg-1(it25)* mutation at the semi-permissive temperature would yield higher levels of centrosome duplication failures and increased embryonic lethality than what each mutant produces on its own.

**Figure 3.**
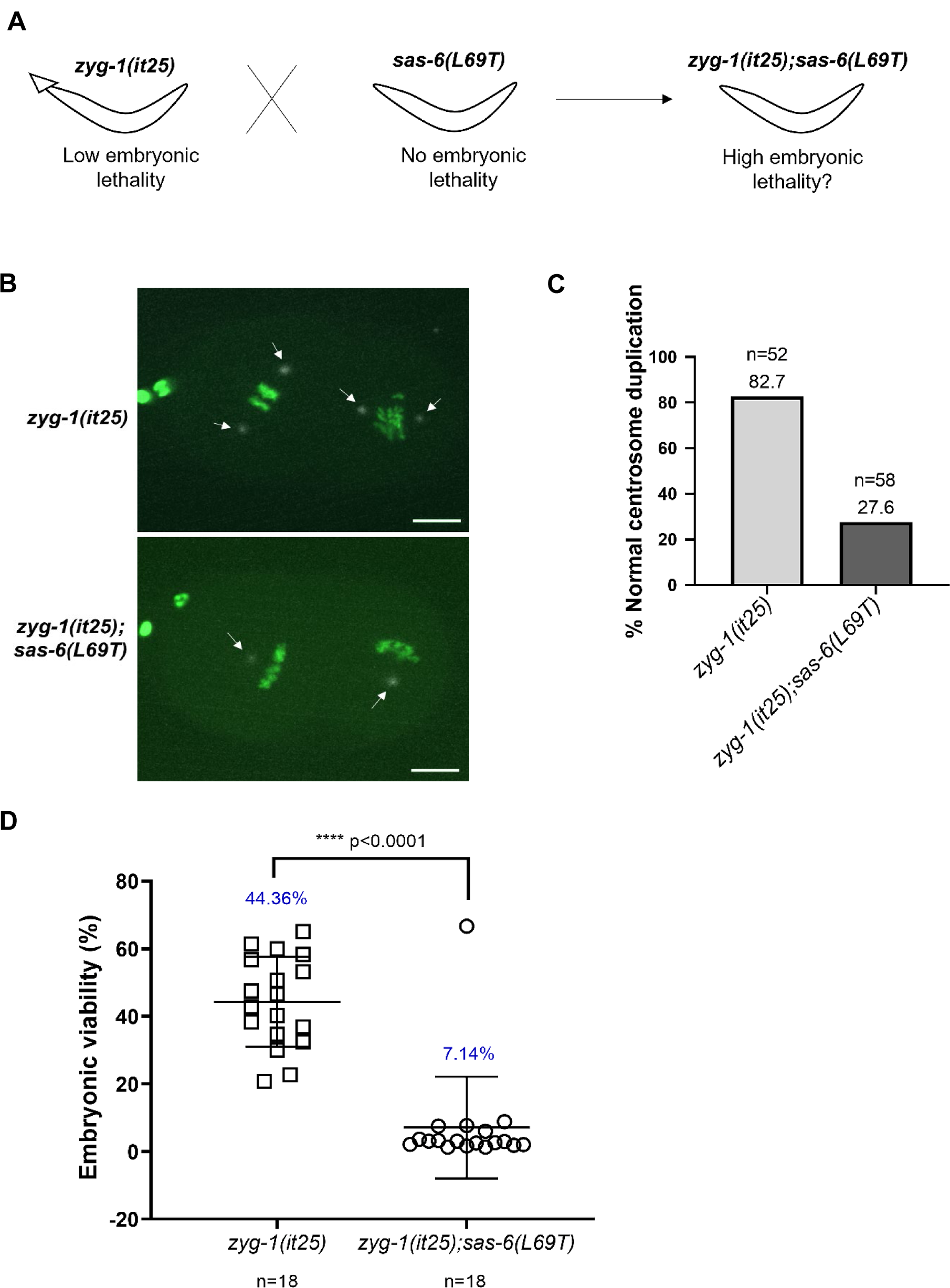
The *sas-6(L69T)* mutation increases the centrosome duplication failures and the embryonic lethality of *zyg-1(it25)* worms. A) Schematic showing the genetic cross that was performed to combine the *sas-6(L69T)* mutation with the *zyg-1(it25)* mutation. B) Stills from time-lapse imaging of *zyg-1(it25)* and *zyg-1(it25);sas-6(L69T)* strains expressing fluorescently-tagged centrosome and DNA markers. Grayscale: mCherry-SPD-2 (centrosome marker); Green: GFP-histone (DNA marker). In a majority of *zyg-1(it25)* mutant embryos, centrosome duplication proceeds normally at a semi-permissive temperature of 21.8°C, yielding two centrosomes in each cell of a 2-cell *C. elegans* embryo (Figure 3B, **top panel, white arrows**). Scale bar = 10 µm. On the contrary, most of the *zyg-1(it25);sas-6(L69T)* mutant centrosomes fail to duplicate at 21.8°C resulting in a single centrosome present in each cell of a 2-cell *C. elegans* embryo (Figure 3B, **bottom panel, white arrows**). C) Quantification of B). n= number of centrosomes analyzed. Approximately 83% of the *zyg-1(it25)* centrosomes duplicate normally at 21.8°C. In contrast, only about 28% of the *zyg-1(it25);sas-6(L69T)* centrosomes duplicate normally at this temperature. D) Quantification of embryonic viability of *zyg-1(it25)* single and *zyg-1(it25);sas-6(L69T)* double mutants at the semi-permissive temperature of 21.7°C. Average embryonic viability is reduced from ∼44% to ∼7% in the presence of the *sas-6(L69T)* mutation. The viability of 1103 embryos from *zyg-1(it25)* worms and 912 embryos from *zyg-1(it25);sas-6(L69T)* worms was analyzed. Embryos from 18 worms were analyzed for each condition. Unpaired two-tailed t-test, p<0.0001. Error bars represent the standard deviation and the middle bar represents the mean.

Upon introducing the *sas-6(L69T)* mutation into the *zyg-1(it25)* strain expressing fluorescent centrosome and chromosome markers, centrosome duplication in the *zyg-1(it25)* and *zyg-1(it25);sas-6(L69T)* worms was assessed at the semi-permissive temperature of 21.8°C **(Figures 3B and 3C)**. Notably, introducing the *sas-6(L69T)* mutation greatly increased the centrosome duplication failures exhibited by the *zyg-1(it25)* mutants at 21.8°C **(Figure 3C)**. Specifically, in the *zyg-1(it25)* single mutant alone, about 83% of the centrosomes analyzed duplicated normally at the semi-permissive temperature of 21.8°C (n=52). However, interestingly, in the *zyg-1(it25);sas-6(L69T)* double mutant, proper centrosome duplication was reduced to only about 28% (n=58). Since accurate centrosome duplication is required for embryonic viability, we next assessed whether embryonic viability of the *zyg-1(it25)* strain was also affected in the presence of the *sas-6(L69T)* mutation.

The embryonic viability of the *zyg-1(it25)* strain expressing fluorescent markers alone at the semi-permissive temperature of 21.7°C was about 44% (number of worms=18; number of progeny= 1103). However, introducing the *sas-6(L69T)* mutation into this background reduced embryonic viability to only about 7% (number of worms=18; number of progeny= 912; unpaired two-tailed t-test p<0.0001) **(Figure 3D)**. These data collectively indicate that the *sas-6(L69T)* mutation mildly inhibits SAS-6 function in regulating centrosome duplication, thereby resulting in a failure of proper centrosome duplication and increased embryonic lethality in the *zyg-1(it25)* background.

### Modeling SAS-6(L69T) using AlphaFold2 suggests that the leucine to threonine substitution at amino acid 69 does not have a major effect on SAS-6 protein structure

The L69T mutation is located within the conserved PISA (Present In SAS-6) motif of SAS-6 **(Figure 4A)**. Further, this amino acid does not have a known involvement in the dimerization of SAS-6 coiled-coil region, in the interaction of SAS-6 N-terminal head domains, in the interaction of SAS-6 with ZYG-1 or in the interaction of SAS-6 with SAS-5 (van Breugel et al. 2011, Kitagawa et al. 2011, Qiao et al. 2013, Hilbert et al. 2013, Lettman et al. 2013, Busch et al. 2019) **(Figure 4A)**. To elucidate the possible mechanism by which the *sas-6(L69T)* mutation affects centrosome duplication, we used AlphaFold2 to predict how the L69T mutation could alter the structure of the SAS-6 protein **(Figures 4B, 4C)**. **Figures 4B and 4C** show the comparison of the SAS-6 PDB model 4G79 from Hilbert et al. 2013 **(Figure 4B)** compared with the AlphaFold2-derived SAS-6(L69T) model **(Figure 4C)** using the ChimeraX program. Our analysis indicates that the overall structural differences due to the L69T mutation in SAS-6 are minor. Therefore, we propose that centriole architecture may not be significantly affected in the *sas-6(L69T)* mutant worms. This is supported by our data that *C. elegans* carrying homozygous copies of the *sas-6(L69T)* mutation are close to 100% viable at all temperatures **(Figure 2B)**. However, the ChimeraX hydrophobicity surface model suggests that a deep hydrophobic pocket that is found within the SAS-6 protein is reduced due to the MCPH-associated L69T mutation **(Figures 4B, 4C, dark blue arrows)**. The function of this hydrophobic pocket within SAS-6 has not yet been elucidated.

**Figure 4.**
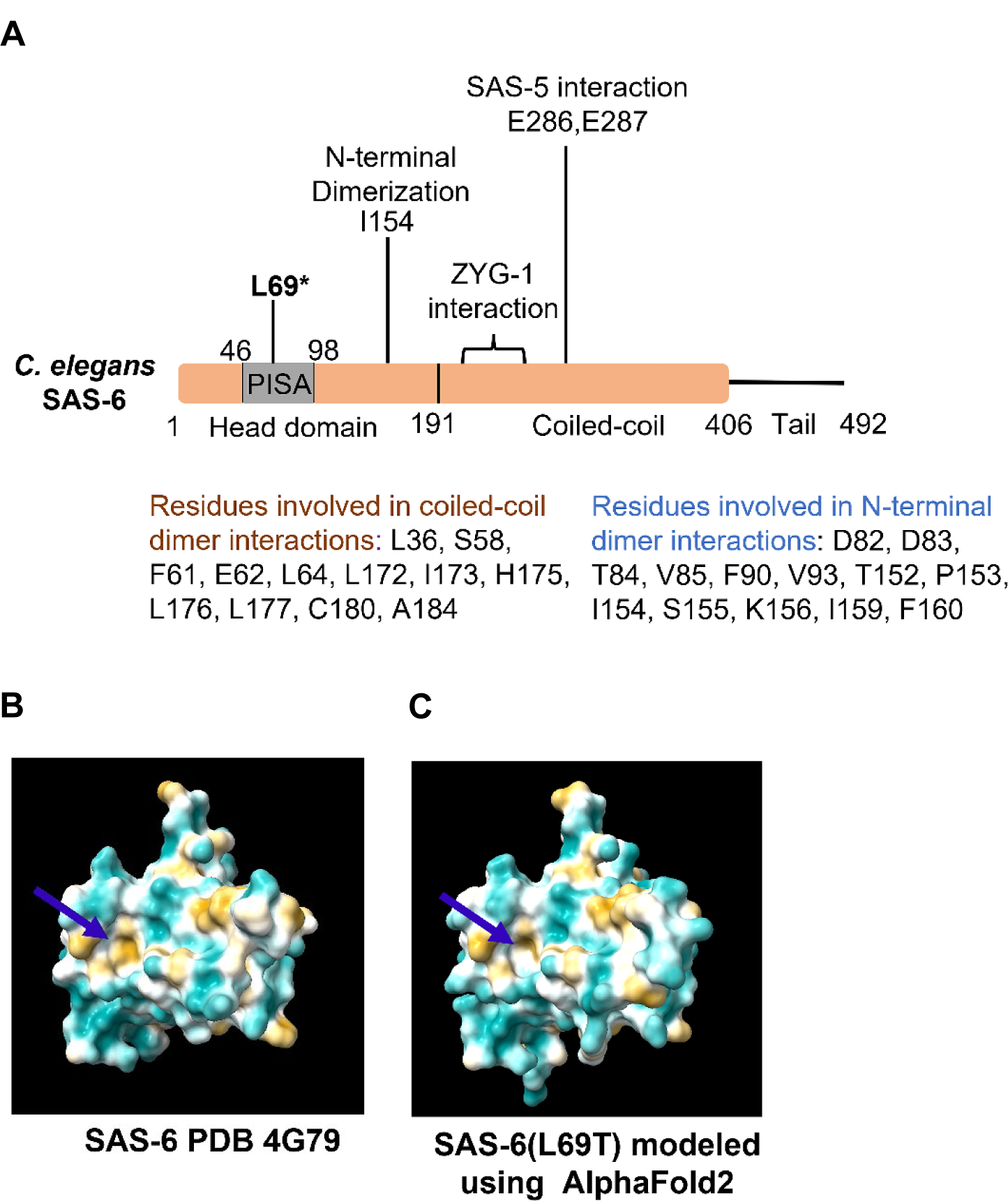
The L69T mutation does not significantly alter the overall structure of the *C. elegans* SAS-6 protein. A) Cartoon depicting the location of the L69T mutation within the *C. elegans* SAS-6 protein oriented with respect to known SAS-6 protein domains and SAS-6 amino acid residues that are important for the interaction of SAS-6 with either itself or with other known interactors (Kitagawa et al. 2011, Lettman et al. 2013). B) Structure of *C. elegans* SAS-6 protein (PDB accession number: 4G79, Hilbert et al. 2013) C) The *C. elegans* SAS-6(L69T) protein structure was modeled using AlphaFold2. The coloring for the models are based on the ChimeraX default palette-option where lipophilicity ranges from −20 to 20 going from cyan (least lipophilic) to golden (most lipophilic). Blue arrows: A deep hydrophobic pocket within the mutant SAS-6(L69T) protein is decreased by the L69T mutation.

### SAS-6 protein levels are altered in the presence of the *sas-6(L69T)* mutation in a *zyg-1(it25)* background but not in a wild-type background

Some of the other known MCPH-associated mutations in centrosome duplication genes such as STIL and PLK4 have been reported to affect their protein levels (Patwardhan et al. 2018, Martin et al. 2014). The effect of *sas-6(L69T)* mutation on SAS-6 protein levels is unknown. Therefore, we questioned whether the *sas-6(L69T)* mutation has any effect on SAS-6 protein levels. To address this, western blotting was performed on whole worm lysates from *zyg-1(it25);sas-6::ha* (control) vs *zyg-1(it25);sas-6(L69T)::ha* worms and from *sas-6::ha* (control) vs *sas-6(L69T)::ha* worms. Western blot analysis indicated that SAS-6 levels decrease on average by 15% in the presence of the *sas-6(L69T)* mutation in the *zyg-1(it25)* background (n=4; p=0.0475, unpaired two-tailed t-test) **(Figure S1A, S1B)**. However, the decrease in SAS-6 protein levels in the presence of the *sas-6(L69T)* mutation in an otherwise wild-type background was not found to be statistically significant (n=4; p=0.1398, unpaired two-tailed t-test) **(Figures S1C, S1D)**.

### Ciliogenesis is perturbed in *sas-6(L69T)* mutant *C. elegans*

Ciliogenesis defects have been linked with the incidence of microcephaly (Alcantara et al. 2014, Gabriel et al. 2016, Broix et al. 2018, Wambach et al. 2018, Zhang et al. 2019, Ding et al. 2019, Sohayeb et al. 2020, Farooq et al. 2020). Defective ciliogenesis is thought to result in cell cycle arrest in neural progenitors during brain development. The loss of neural progenitors due to either cell-cycle arrest or apoptosis has been proposed to be an underlying cause for the incidence of MCPH (Phan and Holland 2021). The effect of the *sas-6(L69T)* mutation on ciliogenesis is currently unknown. Since ciliogenesis defects have been previously linked with MCPH, we questioned whether worms carrying the *sas-6(L69T)* mutation exhibit any ciliogenesis defects. Preliminary dye-filling assays (performed according to the protocol by Power et al. 2020) did not reveal any severe dye-filling defects between control (wild-type N2 worms with HA-tagged endogenous SAS-6) and *sas-6(L69T)* worms. Specifically, we did not observe any worm that completely excluded DiI in their phasmid neurons in either control or *sas-6(L69T)* worms upon dye-filling. Our preliminary experiments revealed that 100% of the control (n=23) and *sas-6(L69T)* (n=25) worms exhibited dye-filling with DiI. Nevertheless, although dye-filling assays are a quick and easy method to visualize cilia defects, a lack of a dye-filling defect phenotype does not exclude the presence of ciliary defects.

To directly determine whether the *sas-6(L69T)* strain exhibits ciliary defects, we obtained a PY6100 strain that expresses a pan-ciliary marker. This strain expresses a GFP transcriptional reporter for OSM-6, a protein that is critical for proper ciliogenesis (Perkins et al. 1986, Collet et al. 1998). Thus, the introduction of *osm-6p::GFP* marks *C. elegans* ciliated sensory neurons with soluble GFP driven from the *osm-6* promoter, allowing for easy visualization of the cilia (Bayer et al. 2020, Xie et al. 2022).

We introduced the *sas-6(L69T)* mutation into the PY6100 strain (*oyIs59 [osm-6p::gfp] III*) by genetic crossing. The resultant strain was named IYR027. The experimental design for combining *osm-6p::GFP* with the *sas-6(L69T)* mutation is shown in **Figure 5A.** The introduction of *osm-6p::GFP* into the *sas-6(L69T)* strain by genetic crossing was confirmed by both confocal microscopy and by genotyping using PCR. For ease of understanding, this new IYR027 strain has been referred to as the *osm-6p::GFP;sas-6(L69T)* strain throughout the paper and has been used to examine the cilia and dendritic defects that arise as a consequence of the *sas-6(L69T)* mutation in **Figures 5, 6 and 7**.

**Figure 5.**
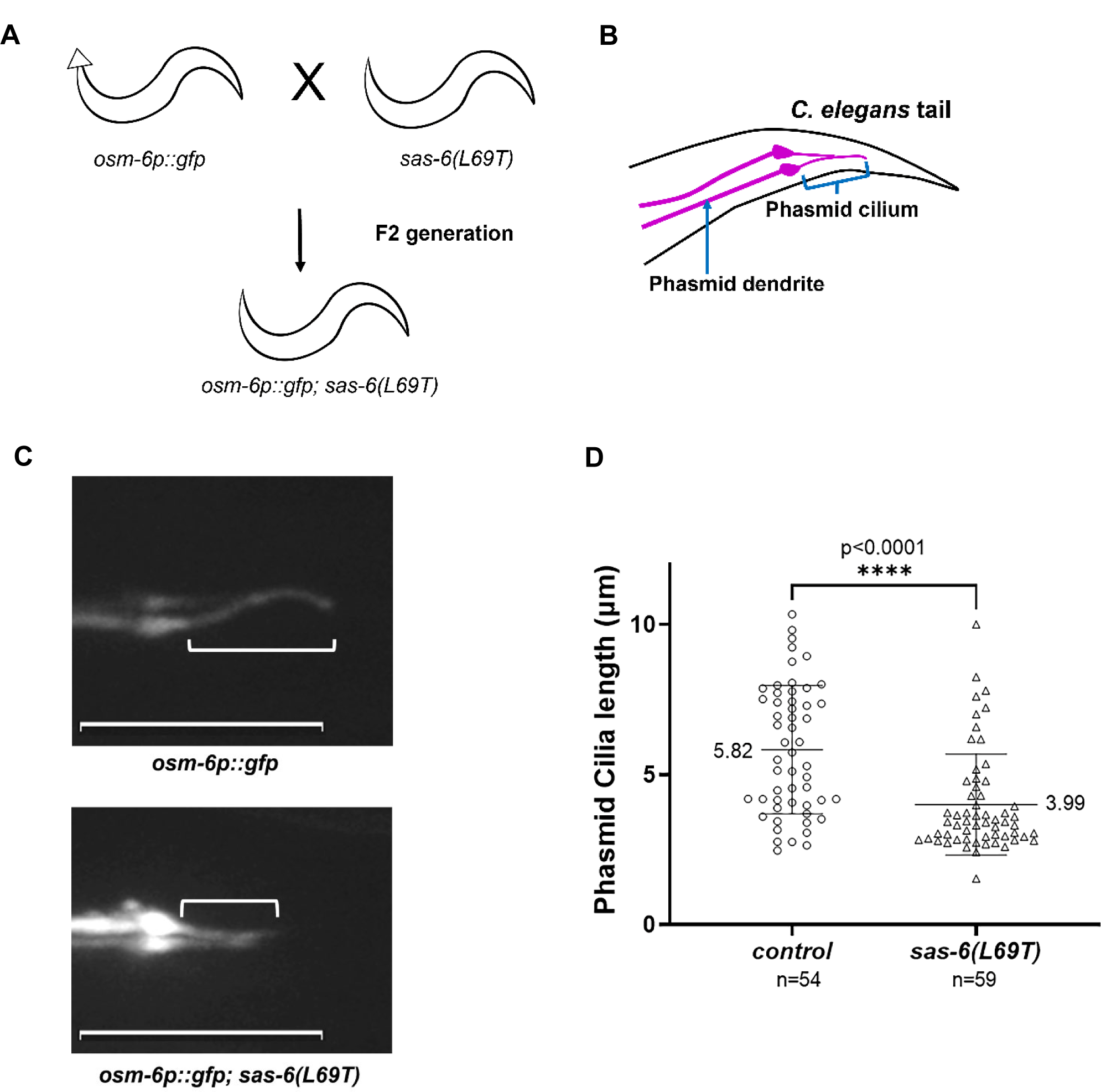
Phasmid cilia length is shorter in *sas-6(L69T)* mutant worms. A) A schematic of the genetic crossing experiment that was performed to introduce the MCPH-associated *sas-6(L69T)* mutation into the PY6100 pan-ciliary marker strain. B) Cartoon showing the anatomical positioning of a *C. elegans* phasmid cilium and dendrite within the tail region of the worm. C) Z-projection confocal images of phasmid neurons from *osm-6p::gfp* (top panel) and *osm-6p::gfp; sas-6(L69T)* (bottom panel) worms highlighting the difference in phasmid cilia lengths between the two genotypes. On average, control *osm-6p::gfp* worms have longer phasmid cilia (top panel, white bracket) than *osm-6p::gfp;sas-6(L69T)* worms (bottom panel, white bracket). White brackets denote the cilia. Scale bar = 10 µm. D) Quantification of phasmid cilia lengths of control (*osm-6p::gfp*) and *sas-6(L69T)* (*osm-6p::gfp;sas-6(L69T)*) worms. Each circle represents the length of a single phasmid cilium. n= number of phasmid cilia analyzed for each genotype. Phasmid cilia length was reduced from 5.82 µm in control worms (n=54) to 3.99 µm in *osm-6p::gfp;sas-6(L69T)* worms (n=59). p<0.0001, unpaired two-tailed t-test. The error bars represent the standard deviation and the middle bar represents the mean.

**Figure 6.**
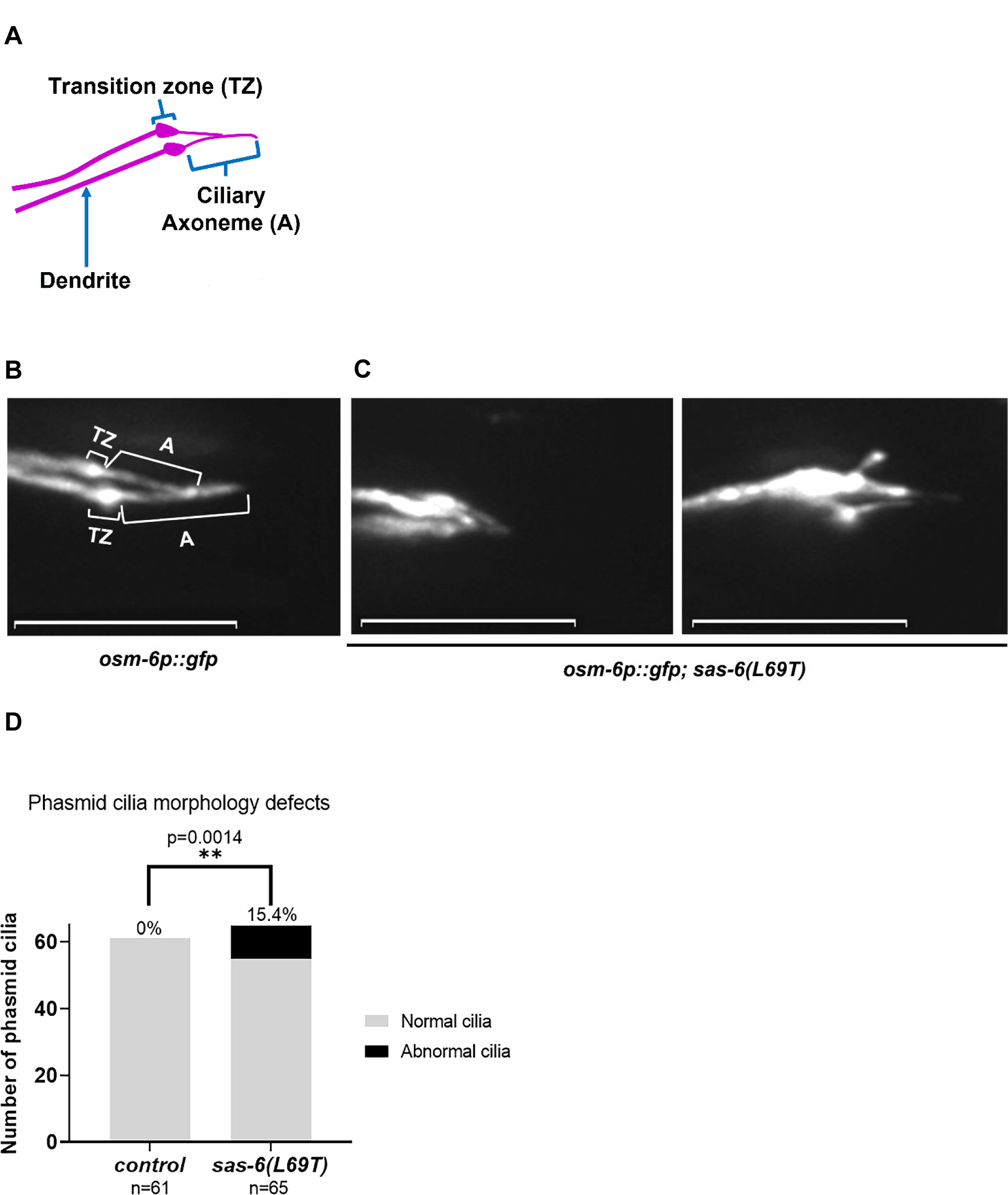
Phasmid cilia morphology is defective in a subset of the *sas-6(L69T)* mutant worms. A) Cartoon showing the organization of a typical phasmid neuron with a ciliary axoneme (A) built on top of the transition zone (TZ). The dendrite of the phasmid neuron connects to the TZ. B) and C) Representative Z-projection confocal images of phasmid cilia morphology in *osm-6p::gfp* and *osm-6p::gfp;sas-6(L69T)* worms, respectively. B) *osm-6p::gfp* worms have a stereotypical phasmid cilia morphology with two prominent TZs at the base and ciliary axonemes projecting from the TZs. Scale bar = 10 µm. C) A subset of the *osm-6p::gfp; sas-6(L69T)* worms exhibit deformed phasmid cilia morphologies. Scale bar = 10 µm. D) Quantification of B) and C). While all of the control (*osm-6p::gfp)* phasmid neurons (n=61) exhibit a normal phasmid cilia morphology, 15.4% of the analyzed *osm-6p::gfp;sas-6(L69T)* phasmid neurons show a deformed cilia morphology (n=65). n= number of phasmid neurons analyzed. p=0.0014, Fisher’s exact test.

**Figure 7.**
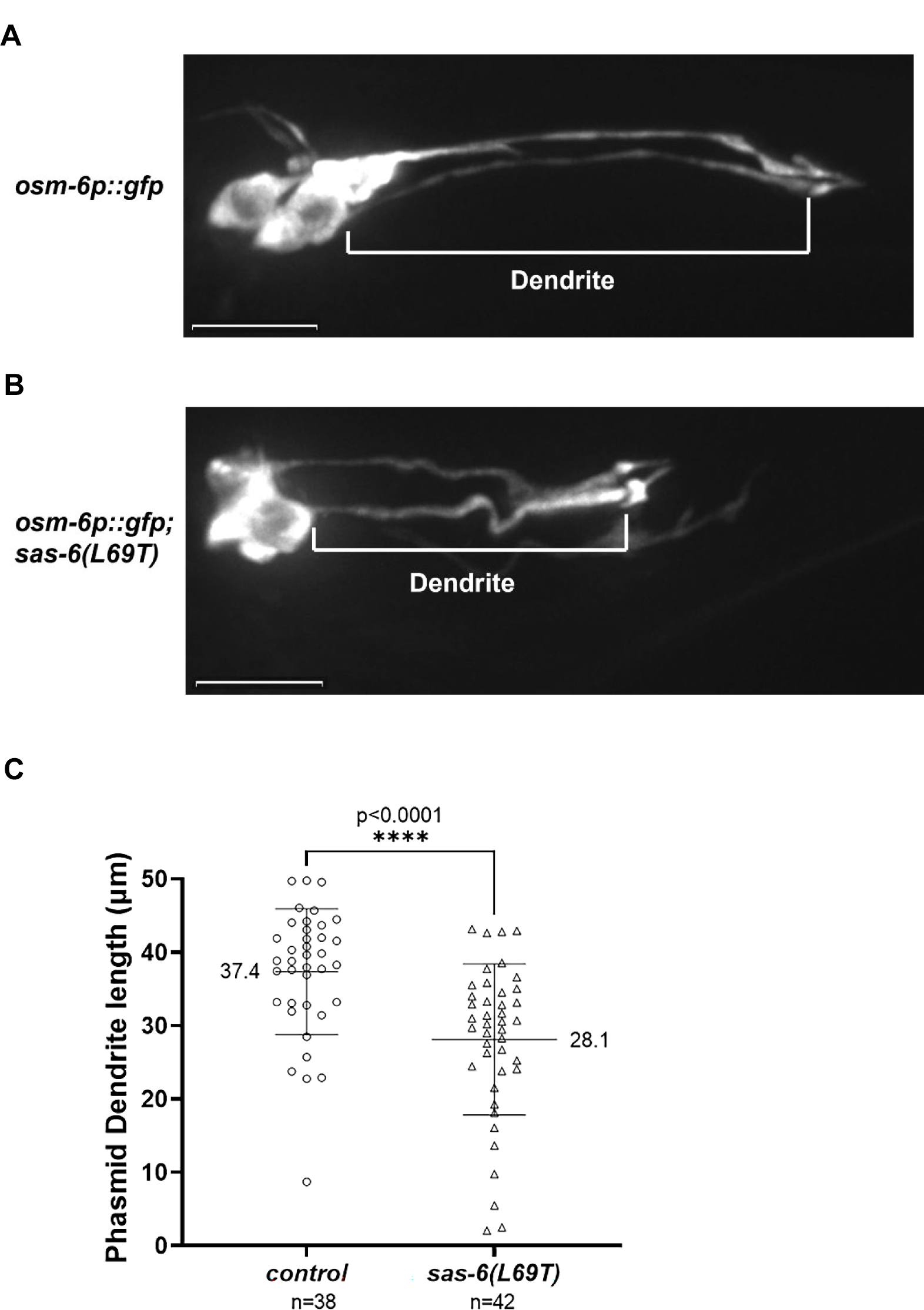
Dendrite length is shorter in *osm-6p::gfp;sas-6(L69T)* phasmid neurons. A) Representative Z-projection confocal image of the average phasmid dendrite length in *osm-6p::gfp* worms. Scale bar = 10 µm. B) Representative Z-projection confocal image of the average phasmid dendrite length in *osm-6p::gfp;sas-6(L69T)* worms. Scale bar = 10 µm. C) Quantification of A) and B). The average dendrite length of control (*osm-6p::GFP*) worms is 37.4 µm (n=38) while that of *osm-6p::gfp;sas-6(L69T)* worms is 28.1 µm (n=42). n= number of phasmid dendrites analyzed. p<0.0001, unpaired two-tailed t-test. The error bars and middle bar represent the standard deviation and the mean, respectively.

Since the *sas-6(L69T)* strain also has an HA tag on the mutant SAS-6 protein, to create an appropriate control strain, we first crossed the PY6100 strain with a strain expressing SAS-6::HA to account for the potential effect of the HA-tag on SAS-6 function in regulating ciliogenesis. This strain was named IYR026. We then CRISPR-edited the IYR026 strain to make it genetically identical to the *osm-6p::GFP;sas-6(L69T)* strain except for the fact that this new strain lacked the leucine 69 to threonine mutation. For convenience, we refer to this control strain (IYR040) as the *osm-6p::GFP* strain. This strain served as the control for all the cilia and dendrite analysis conducted in **Figures 5, 6 and 7**.

To investigate whether *sas-6(L69T)* mutant worms exhibit defects in ciliogenesis, we examined the cilia of the phasmid neurons which are present in the posterior region of the worms **(Figure 5B)**. Intriguingly, upon measuring the length of the phasmid neuronal cilia, we found a significant decrease in the average ciliary length in the *osm-6p::GFP;sas-6(L69T)* mutant worms **(Figure 5C, bottom panel, white bracket)** as compared with control (*osm-6p::GFP)* worms **(Figure 5C, top panel, white bracket)**. Specifically, control *osm-6p::GFP* worms had an average phasmid cilia length of about 5.82 µm (n=54) while the *osm-6p::GFP; sas-6(L69T)* worms had shorter cilia with an average phasmid cilia length of only 3.99 µm (n=59) (unpaired two-tailed t-test, p<0.0001). These data indicate that the *sas-6(L69T)* mutation indeed perturbs ciliogenesis in *C. elegans*.

### Phasmid cilia morphology and dendrite length are defective in the *sas-6(L69T)* mutant worms

While analyzing cilia length of *osm-6p::GFP;sas-6(L69T)* worms, we found that some of the cilia from this strain exhibited a disfigured cilia morphology as compared with control worms. Our analysis revealed that 100% of the phasmid cilia of control (*osm-6p::gfp)* worms (n=61) showed a normal phasmid cilia morphology with two distinct transition zones and a ciliary axoneme projecting from each transition zone **(Figure 6A, 6B, 6D)**. However, 10 out of 65 (15.4%) of the *osm-6p::GFP;sas-6(L69T)* phasmid cilia exhibited an abnormal morphology **(Figures 6A, 6C, 6D).** This abnormal phasmid cilia morphology defect of the *osm-6p::GFP; sas-6(L69T)* worms was found to be statistically significant via a Fisher’s exact test (p=0.0014). These data indicate that the *sas-6(L69T)* mutation perturbs phasmid cilia morphology in a subset of the phasmid cilia.

In addition to the phasmid cilia morphology defect, the *osm-6p::GFP;sas-6(L69T)* worms also displayed a dendritic extension defect. In general, the average lengths of the phasmid dendrites of control *osm-6p::GFP* worms **(Figure 7A)** were longer than those of *osm-6p::GFP*;*sas-6(L69T)* worms **(Figure 7B)**. Specifically, phasmid dendrites analyzed from control worms exhibited an average length of 37.4 µm (n=38). In contrast, the average length of phasmid dendrites of *sas-6(L69T)* worms was 28.1 µm (n=42) **(Figure 7C).** This phasmid dendrite length defect of the *osm-6p::GFP;sas-6(L69T)* worms was found to be statistically significant via an unpaired two-tailed t-test (p<0.0001).

### Chemotaxis is defective in the *sas-6(L69T)* mutant worms

Since defects in ciliogenesis in the *C. elegans* sensory neurons often affect worm behaviors, we investigated the effect of the *sas-6(L69T)* mutation on chemotaxis to the chemoattractant butanone. The experimental scheme for the chemotaxis assays is shown in **Figure 8A**. Our studies demonstrated that *C. elegans* carrying the *sas-6(L69T)* mutation exhibited a reduced attraction to 1:500 butanone as compared with control worms (two-tailed unpaired t-test, p<0.0001) **(Figures 8B, 8C)**. Specifically, our data showed that the average chemotaxis index of control worms (*sas-6::ha*) was 0.4107 (n=2825). In comparison, the average chemotaxis index of *sas-6(L69T)* worms was reduced to 0.3307 (n=1711) **(Figures 8B, 8C)**. These data indicate that defective ciliogenesis in the *sas-6(L69T)* mutant worms likely causes a defective *C. elegans* chemotaxis response to 1:500 butanone. Hence, SAS-6 function in regulating ciliogenesis is likely to be important for mediating certain worm behaviors such as chemotaxis.

**Figure 8.**
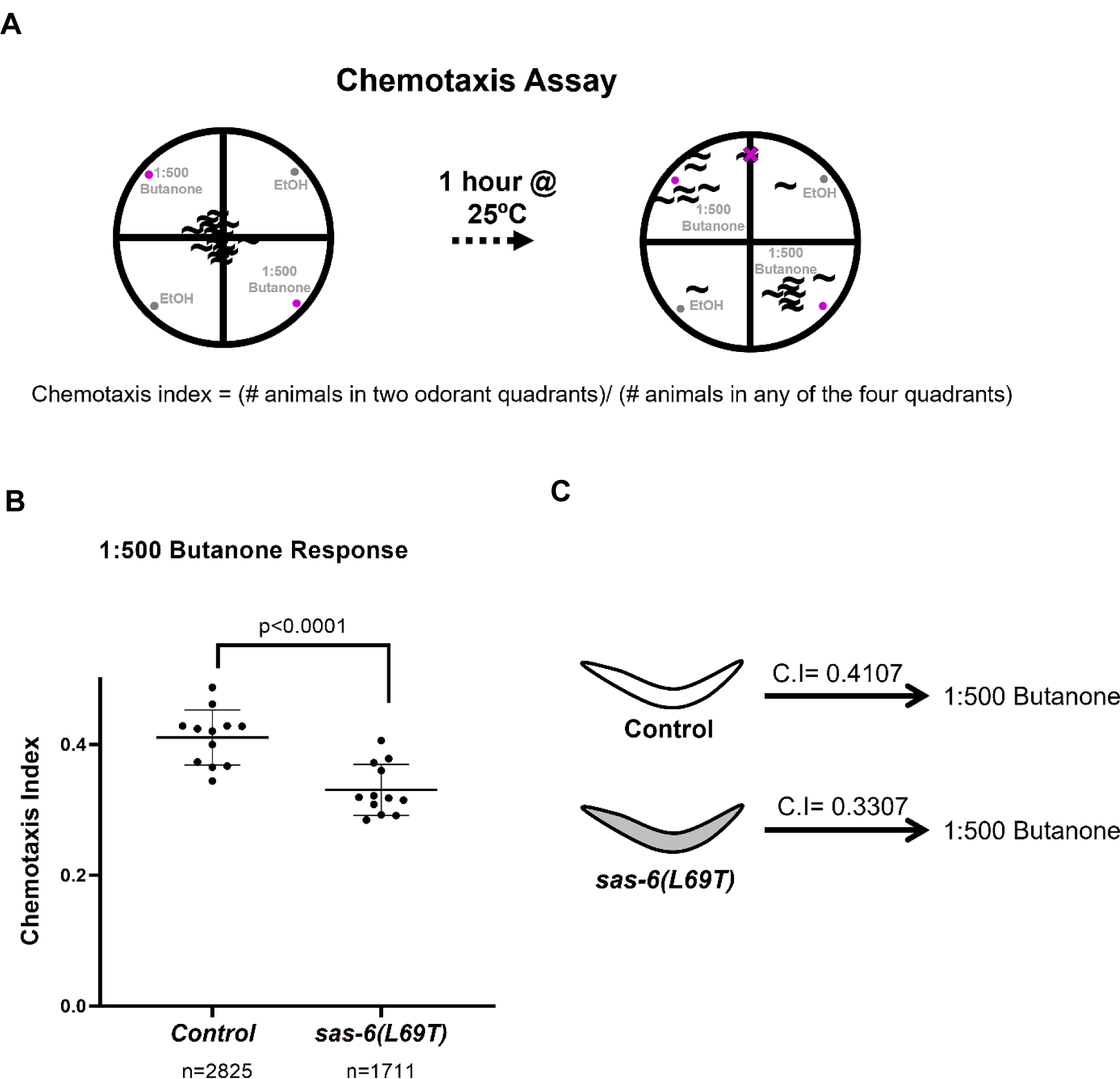
The *sas-6(L69T)* mutation reduces *C. elegans* chemotaxis to 1:500 Butanone. A) Schematic of chemotaxis assay experiments. L4 stage *C. elegans* were plated onto unseeded NGM plates that had been divided into four quadrants, two with solvent (ethanol) and two with odorant (1:500 butanone in ethanol). The worms were placed in the center of the plate and allowed to move around for 1 hour at 25°C. After 1 hour, the number of worms in each quadrant were counted and quantified. The formula C.I. = (# animals in two odorant quadrants)/(# animals in any of the four quadrants) was used to compute the Chemotaxis Index (C.I.). Worms on any line or within the origin were not counted. Four plates were used for each genotype per experiment. Assays were repeated in triplicate. B) Quantification of data obtained from A). The *sas-6(L69T)* mutants exhibit a decreased chemotaxis towards 1:500 butanone as compared with control worms. n= Number of worms whose chemotaxis towards 1:500 butanone was analyzed. p<0.0001, unpaired two-tailed t-test. C) Summary of data from A) The average chemotaxis index of *sas-6(L69T)* worms was 0.3307. This chemotaxis index was less than the average chemotaxis index of control worms which was 0.4107. C.I.=Chemotaxis Index.

Interestingly, chemotaxis towards butanone is predominantly controlled by the AWC amphid sensory neurons (Bargmann et al. 1993). These data indicate that ciliogenesis may also be perturbed in the amphid sensory neurons in *sas-6(L69T)* mutant worms. In the present study, we have mainly focused our efforts on analyzing ciliogenesis in the *C. elegans* phasmid neurons. Future studies should be directed towards determining whether ciliogenesis in the amphid neurons is also affected in *C. elegans* carrying the *sas-6(L69T)* mutation.

## Discussion

Through our studies, we have re-created the MCPH-associated *sas-6(L69T)* mutation in *C. elegans* by CRISPR to generate the first *C. elegans* model of a MCPH-associated mutation. A previous study investigated the effects of the human SASS6(I62T) mutation (the human equivalent of the SAS-6(L69T) mutation) on centrosome duplication in U2OS cells (Khan et al. 2014). In their studies, the authors noted an inhibition of centrosome duplication in U2OS cells expressing the SASS6(I62T) protein from exogenous plasmid constructs carrying the *SASS6(I62T)* mutation. The levels of SASS6 in the cell must be tightly regulated for its proper cellular function. Either overexpressing the SASS6 protein or depleting it can affect proper centrosome duplication (Dammermann et al. 2004, Leidel et al. 2005). Therefore, to investigate the effect of the *sas-6(L69T)* mutation on centrosome duplication in an endogenous context, we monitored centrosome duplication in our *sas-6(L69T)* CRISPR worm model. Interestingly, *C. elegans* carrying this mutation appeared phenotypically wild-type and did not exhibit any defects in embryonic viability or brood size **(Figure 2)**. *C. elegans* embryonic divisions are reliant on proper centrosome duplication. Since we did not detect any embryonic lethality in the presence of the *sas-6(L69T)* mutation **(Figure 2B)**, we concluded that centrosome duplication is likely to be relatively unaffected in the *sas-6(L69T)* mutant worms in a wild-type background.

To uncover the potential milder effects of this mutation on centrosome duplication that may not be visible in a wild-type genetic background, we introduced this mutation in the sensitized *zyg-1(it25)* background. Interestingly, the *sas-6(L69T)* mutation significantly enhanced the centrosome duplication failures and embryonic lethality of *zyg-(it25)* worms **(Figures 3B, 3C, 3D)**. These data demonstrate that the MCPH-associated *sas-6(L69T)* mutation does, in fact, reduce SAS-6 function in regulating centrosome duplication. However, this effect can only be revealed in a sensitized genetic background. Thus, at least in the context of this MCPH-associated mutation, centrosome duplication is only mildly affected by this mutation. Future studies will involve elucidating the molecular mechanism by which the *sas-6(L69T)* mutation inhibits centrosome duplication.

Our data that the *sas-6(L69T)* mutation likely has a mild effect on SAS-6 function in regulating centrosome duplication are consistent with the observed phenotypic effects of the *SASS6(I62T)* mutation in human MCPH patients. *SASS6*/*sas-6* is an essential gene that is required for proper centrosome duplication and cell division (Kamath et al. 2003, Dammermann et al. 2004, Leidel et al. 2005, Blomen et al. 2015). If the *SASS6(I62T)* mutation severely inhibited SASS6 function in regulating centrosome duplication, it would be expected to result in embryonic lethality. As a result, there would not be human patients living with homozygous copies of the *SASS6(I62T)* mutation. However, the fact that there are *SASS(I62T)* homozygous individuals in the population (Khan et al. 2014) indicates that this mutation does not severely perturb SASS6 role in regulating centrosome duplication in humans. Therefore, our data in *C. elegans* which demonstrate that embryonic viability is unaffected in the presence of this mutation **(Figure 2B)** are in line with the observations in human MCPH patients carrying this mutation.

Modeling the L69T mutation using AlphaFold2 did not reveal any significant changes in the structure of the SAS-6 protein **(Figures 4B, 4C)**. This is not a surprising finding since *sas-6* is an essential gene and therefore severely affecting SAS-6 structure may alter centriole architecture and result in embryonic lethality. However, our AlphaFold2 analysis revealed that a deep hydrophobic pocket within SAS-6 is altered by the *sas-6(L69T)* mutation **(Figures 4B, 4C, dark blue arrows)**. Presently, no known function has been ascribed to this hydrophobic pocket in SAS-6. Based upon our data, we are tempted to speculate that this SAS-6 hydrophobic pocket may mediate an as-yet unknown interaction of SAS-6 with a protein(s) that may be important for its function in ciliogenesis, dendrite morphogenesis and/or centrosome duplication.

The effect of the *SASS6(I62T)* mutation on ciliogenesis has not been investigated. Upon introducing the *sas-6(L69T)* mutation into a pan-ciliary marker expressing *C. elegans* strain, we determined that ciliogenesis is perturbed in the *sas-6(L69T)* mutant worms **(Figures 5C, 5D, 6B, 6C, 6D)**. Specifically, the *sas-6(L69T)* worms have shorter phasmid cilia and a defective phasmid cilia morphology **(Figures 5C, 5D, 6B, 6C, 6D)**. Ciliogenesis defects have been associated with the incidence of microcephaly (Alcantara et al. 2014, Gabriel et al. 2016, Broix et al. 2018, Wambach et al. 2018, Zhang et al. 2019, Ding et al. 2019, Sohayeb et al. 2020, Farooq et al. 2020). Interestingly, in a study by Shohayeb et al., the authors determined that mouse embryonic fibroblasts that were derived from WDR62 mutant mice carrying CRISPR-edited MCPH patient mutations exhibited a shortened ciliary length, like we report here (Shohayeb et al. 2019). In another study, Wambach et al. cultured MCPH patient derived fibroblasts and found that these fibroblasts had shorter primary cilia than those of fibroblasts that were cultured from a healthy control individual (Wambach et al. 2018). Thus, there is precedence for shorter ciliary length being associated with MCPH.

Our data indicate that SAS-6 levels are reduced in *sas-6(L69T)* worms in a *zyg-1(it25)* background but not in an otherwise wild-type background **(Figure S1)**. Since we observe ciliogenesis defects in the presence of the *sas-6(L69T)* mutation even in a wild-type background (Figures 5C, 5D, 6B, 6C, 6D), we propose that decreased SAS-6 levels are likely not sufficient to explain the effects of the *sas-6(L69T)* mutation.

We find that in *C. elegans* carrying the *sas-6(L69T)* mutation, centrosome duplication defects can only be detected in a sensitized *zyg-1(it25)* genetic background **(Figures 3B, 3C).** However, the ciliogenesis and dendritic defects induced by this mutation can be uncovered even in an otherwise wild-type background **(Figures 5C, 5D, 6B, 6C, 6D)**. These findings lead us to speculate that perhaps, at least in the context of this specific MCPH-associated mutation, the inhibition of proper ciliogenesis may be a more important contributor to the development of MCPH than the inhibition of centrosome duplication. However, it is also possible that the centrosome duplication and ciliogenesis defects induced by this mutation synergize to cause MCPH in human patients carrying this mutation. In the future, it will be interesting to determine if the *SASS6(I62T)* mutation also affects ciliogenesis in human cell lines. It will also be interesting to examine if other such MCPH-associated mutations likewise affect ciliogenesis in addition to affecting centrosome duplication.

Since *sas-6* is an essential gene, many *sas-6* mutant alleles are difficult to maintain and analyze (O’Rourke et al. 2011, Song et al. 2011, The C. elegans Deletion Mutant Consortium et al. 2012, Lettman et al. 2013). This has made it tedious to dissect the function of SAS-6 in ciliogenesis. A previous study expressed a *sas-6* sgRNA and Cas9 protein using a heat-shock promoter to demonstrate that SAS-6 plays a critical role in *C. elegans* ciliogenesis (Li et al. 2017). However, the mechanistic investigation of these defects has been challenging due to the high lethality and sterility of *sas-6* mutant alleles. Notably, our studies indicate that *sas-6(L69T)* mutant worms appear phenotypically wild-type and yet exhibit significant ciliogenesis defects. Therefore, we expect that our *sas-6(L69T)* CRISPR strain will be a useful tool to unravel the mechanism by which SAS-6 regulates ciliogenesis. In the future, forward genetic screens can be performed with the *sas-6(L69T)* allele to identify additional genes that may genetically interact with *sas-6* to regulate ciliogenesis in the worm.

Our data indicate that *C. elegans* harboring the *sas-6(L69T)* mutation also exhibit sensory neuronal defects such as a shortened dendrite length **(Figure 7)**. This is a surprising finding since *C. elegans* centrioles have been found to degenerate during neuronal differentiation (Li et al. 2017, Serwas et al. 2017, Nechipurenko et al. 2017). One possibility is that SAS-6 functions independently of its role as a core centriolar protein in regulating dendrite length in *C. elegans* sensory neurons. Another possibility is that SAS-6 function at the centrioles may be required for proper phasmid dendrite extension through an unknown mechanism. In the future, transmission electron microscopy analysis of dendrites and cilia from *sas-6(L69T)* mutant worms should be performed to determine the nature of the dendritic and ciliary defects in these mutant worms. Future studies will also involve analyzing *C. elegans* strains expressing a variety of fluorescently-tagged markers (e.g. Intraflagellar transport markers, transition zone markers, neuron-specific markers, etc.) to further dissect the mechanism by which the *sas-6(L69T)* mutation affects ciliogenesis and dendrite extension.

Defective chemotaxis towards butanone was seen in *C. elegans* carrying the *sas-6(L69T)* mutation **(Figure 8).** Future studies should also be directed towards examining what other worm behaviors (if any) are altered in *C. elegans* carrying the *sas-6(L69T)* mutation. It is important to note that in this study, we have only examined the effect of the *sas-6(L69T)* mutation on ciliogenesis and behaviors in the *C. elegans* hermaphrodite. The effect of this mutation on *C. elegans* males is unknown. Assays such as male mating assays could be carried out in the future to determine if male-specific behaviors are affected by the *sas-6(L69T)* mutation.

As our *C. elegans* model of the *sas-6(L69T)* mutation recapitulates many of the effects seen in the human MCPH patients carrying this mutation, we expect that *C. elegans* will be more widely adopted as a model system to study the effects of additional MCPH-associated human mutations in the future.

## Data Availability Statement

All the strains made and used in this study will be made available upon request. **Table S1** lists all the strains made in this study.

## Acknowledgments

We would like to thank the Sengupta lab for providing the PY6100 strain that was used for cilia analysis. We would like to thank the O’Connell lab for providing the OC14 and OC869 strains that have been used in this study. The wild-type N2 Bristol strain was provided by the CGC, which is funded by NIH Office of Research Infrastructure Programs (P40 OD010440). We would like to acknowledge Peter Nunan and Mariah Moreland for their technical assistance. Adam Norris received funding from NIH R35GM133461. Finally, we would like to thank Drs. Steve Caplan and Nina Peel for their critical comments on this manuscript.

---

**Figure S1.**
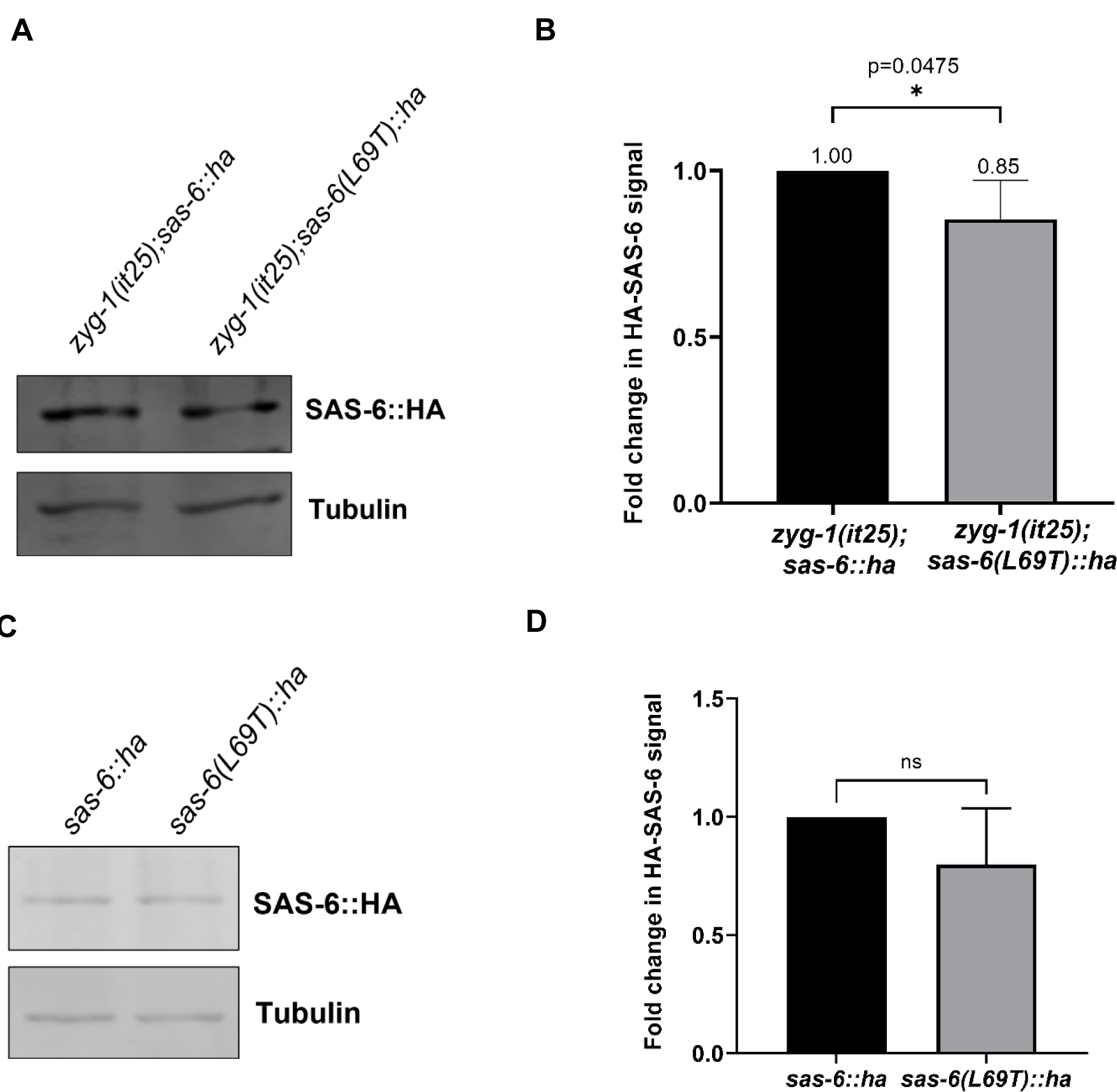
SAS-6 protein levels are slightly reduced in the presence of the *sas-6(L69T)* mutation in a *zyg-1(it25)* background but are unaffected in a wild-type background. A) A representative western blot image of an HA blot showing SAS-6 levels in *zyg-1(it25);sas-6::ha* and *zyg-1(iti25);sas-6(L69T)::ha* worms. Tubulin served as a loading control. SAS-6 protein levels are slightly decreased in the presence of the *sas-6(L69T)* mutation in a *zyg-1(it25)* background B) Quantification of A) from four independent experiments. On average, SAS-6 protein levels are reduced by 15% in the presence of the *sas-6(L69T)* mutation in the *zyg-1(it25)* background. P=0.0475, unpaired two-tailed t-test. C) A representative western blot image of a HA blot showing SAS-6 levels in *sas-6::ha* and *sas-6(L69T)::ha* worms. Tubulin was used as a loading control. Although SAS-6 protein levels tend to decrease in the presence of the *sas-6(L69T)* mutation in a wild-type background, this decrease is not statistically significant (p=0.1398, unpaired two-tailed t-test).

