## Supplementary figures and images for "A primary microcephaly-associated *sas-6* mutation perturbs centrosome duplication, dendrite morphogenesis, and ciliogenesis in *Caenorhabditis elegans*"

### Figure S1

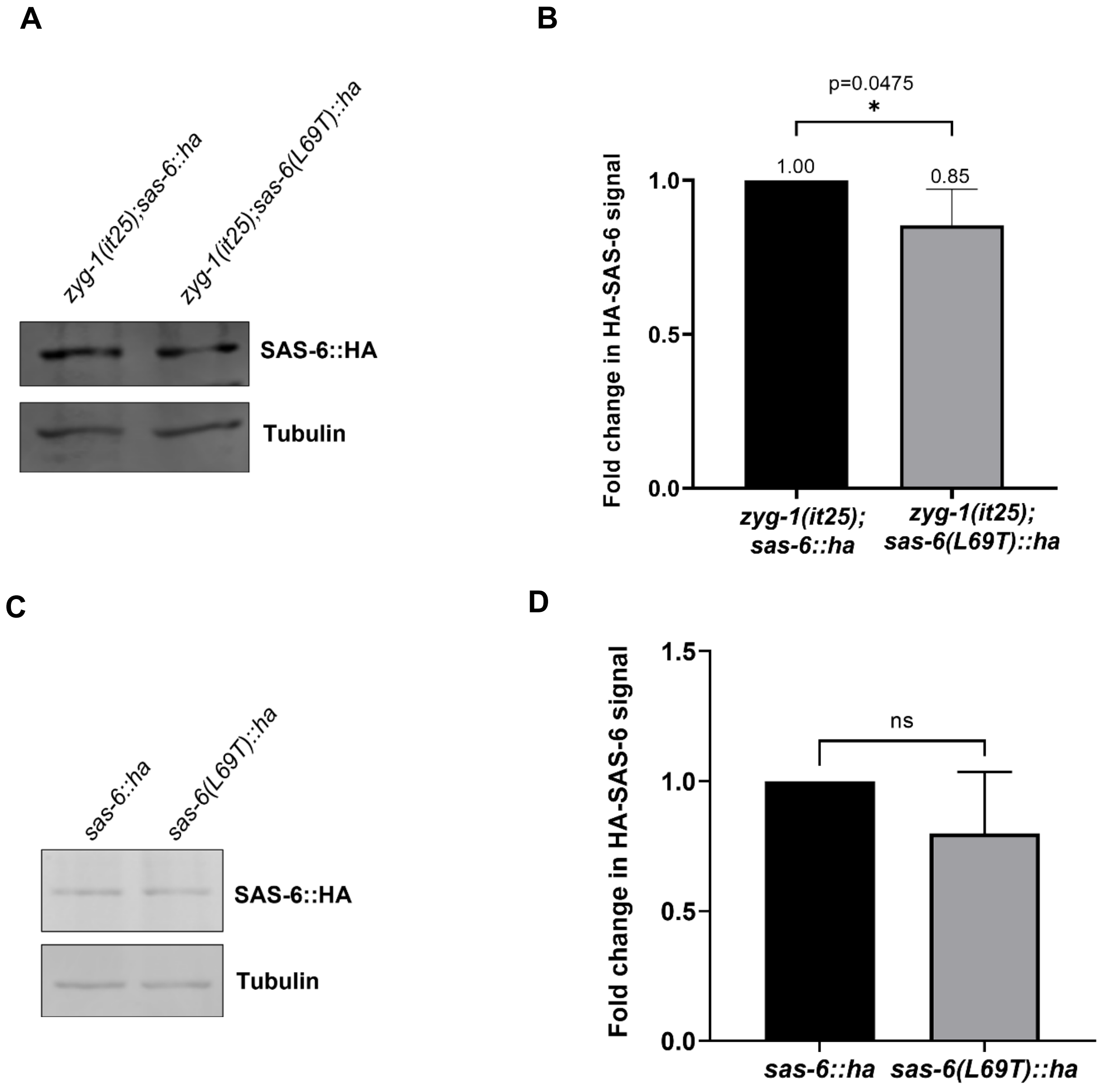
