## Supplementary material for "A primary microcephaly-associated *sas-6* mutation perturbs centrosome duplication, dendrite morphogenesis, and ciliogenesis in *Caenorhabditis elegans*": Tables S1-S3

**Table S1. *C. elegans* strains used in this study**

| STRAIN | GENOTYPE | METHOD | RESOURCE |
|---|---|---|---|
| N2 | wild-type | N/A | CGC |
| PY6100 | *oyIs59 [osm-6p::gfp] III* | N/A | Piali Sengupta Lab |
| OC14 | *zyg-1(it25) II* | N/A | Kevin O'Connell Lab |
| OC869 | *bsSi15 [pKO109: spd-2p::spd-2::mCherry::spd-2_3'UTR, unc-119(+)] I; zyg-1(it25) bsSi30 [pCW9: unc-119(+) pcdk-11.2::sfgfp::his-58::cdk-11.2_3' UTR] II* | N/A | Kevin O'Connell Lab |
| IYR001 | *sas-6(luv1[sas-6::ha]) IV* | CRISPR | Bergwell *et al.* 2019 |
| IYR002 | *sas-6(luv2[(sas-6[L69T]::ha)]) IV* | CRISPR | This study |
| IYR003 | *zyg-1(it25) II; sas-6(luv1[sas-6::ha]) IV* | Crossed IYR001 with OC14 | This study |
| IYR004 | *zyg-1(it25) II; sas-6(luv2[(sas-6[L69T]::ha)]) IV* | Crossed IYR002 with OC14 | This study |
| IYR005 | *bsSi15 [pKO109: spd-2p::spd-2::mCherry::spd-2_3'UTR, unc-119(+)] I; zyg-1(it25) bsSi30 [pCW9: unc-119(+) pcdk-11.2::sfgfp::his-58::cdk-11.2_3' UTR] II; sas-6(luv1[sas-6::ha]) IV* | Crossed IYR001 with OC869 | This study |
| IYR006 | *bsSi15 [pKO109: spd-2p::spd-2::mCherry::spd-2_3'UTR, unc-119(+)] I; zyg-1(it25) bsSi30 [pCW9: unc-119(+) pcdk-11.2::sfgfp::his-58::cdk-11.2_3' UTR] II; sas-6(luv2[(sas-6[L69T]::ha)]) IV* | Crossed IYR002 with OC869 | This study |
| IYR026 | *oyIs59 [osm-6p::gfp] III; sas-6(luv1[sas-6::ha]) IV* | Crossed IYR001 with PY6100 | This study |
| IYR027 | *oyIs59 [osm-6p::gfp] III; sas-6(luv2[(sas-6[L69T]::ha)]) IV* | Crossed IYR002 with PY6100 | This study |
| IYR040 | *oyIs59 [osm-6p::gfp] III; sas-6(luv40) IV* | CRISPR-edited IYR026 to make it genetically identical to IYR027 except that it lacks the L69T mutation | This study |

**Table S2. Sequences of guide RNAs, repair templates and primers used for CRISPR/Cas9 genome editing in this study**

| Gene | Type of oligo | Sequence (5' to 3') |
|---|---|---|
| ***dpy-10*** (Paix et al. 2015) | co-CRISPR crRNA | GCUACCAUAGGCACCACGAG |
| ***sas-6*** (Bergwell et al. 2019) | C-terminal HA-tag crRNA | AUUUUAUCGUUGAGCGGGUG |
| ***sas-6*** | crRNA used to introduce the L69T mutation and to make the control IYR040 strain | UGAUUUUGAAUUUCUAUUCU |
| ***dpy-10*** (Arribere et al. 2014) | *dyp-10(cn64)* repair oligo | CACTTGAACTTCAATACGGCAAGATGAGAATGAC TGGAAACCGTACCGCATGCGGTGCCTATGGTAG CGGAGCTTCACATGGCTTCAGACCAACAGCCTAT |
| ***sas-6*** (Bergwell et al. 2019) | HA repair oligo | TATTTTCAAGTAAAGGACAAGAAAAAATCAATAAAA AAGATTTTAAGCGTAATCTGGAACATCATATGGGTA ACGCTGTGCCGGCGGAGTGTTCGTCACACTTGAA CCAGTAGTCTCGTCGGCGATTAGTTGA |
| ***sas-6*** | L69T repair oligo | AAAGGAGCTAAAATTCGAAATCAGCCGCTCCGAT GATTTTGAATTCCTCTTTTCAGAGACAACG AACAACGAGAAATATCAGATTTTGGCTCGAGATCA CGATTTAACA |
| ***sas-6*** | Control repair oligo used to make IYR040 | AAAGGAGCTAAAATTCGAAATCAGCCGCTCCGAT GATTTTGAATTCCTCTTTTCAGAGACATTG AACAACGAGAAATATCAGATTTTGGCTCGAGATCA CGATTTAACA |
| ***sas-6*** | Primers for *sas-6(luv2)* and *sas-6(luv40)* screening | Forward primer: GGAACGAAACCAGTGGAGAA<br>Reverse primer: GAAATAGGAGTCTTCGAGAA |
| ***sas-6*** (Bergwell et al. 2019) | Primers for *sas-6(luv1)* screening (Bergwell et al. 2019) | Forward primer: CCCCATTCCGTGACAATACA<br>Reverse primer: CCTTACCTCTTGAACTGCC |

**Table S3. Antibodies used for western blotting**

| Antibodies | Host | Manufacturer | Catalog# | Application | Dilution |
|---|---|---|---|---|---|
| Alpha tubulin | Mouse | Santa Cruz Biotechnology | sc-32293 | IB | 1:200 |
| HA | Rabbit | Cell Signaling | 3724S | IB | 1:1000 |
| Mouse IRDye 680 RD | Goat | LI-COR | 926-68070 | IB | 1:14000 |
| Rabbit IRDye 800CW | Donkey | LI-COR | 926-32213 | IB | 1:14000 |
